# Longitudinal Associations Between Screen Time, Brain Development, and Language Outcomes in Early Childhood

**DOI:** 10.1101/2025.08.25.672107

**Authors:** Eun Jung Choi, Emily S. Nichols, Lianne Tomfohr-Madsen, Gerald F. Giesbrecht, Kathryn Y. Manning, Catherine A. Lebel, Emma G. Duerden

**Author notes:** **Address correspondence to:** Emma G. Duerden, PhD, Applied Psychology, Faculty of Education, 1137 Western Rd, London, Ontario N6G 1G7.

## Abstract

Language development during toddlerhood is supported by both neurobiological maturation and environmental experiences. It relies on reciprocal interaction, and excessive screen time exposure may have a negative impact. In the current study, we investigated how screen time at age two relates to language outcomes and brain development at ages two and three. Seventy toddlers underwent MRI scanning and neurodevelopmental testing, and brain volumes in language-related areas were extracted. Structural equation modelling showed that at age two, there was a negative relationship between screen time and pars triangularis volumes. Importantly, smaller volumes at age two predicted greater screen time usage at age three, mediated by poorer language outcomes. These results suggest that over time, children with smaller volumes and weaker language skills at age 2 became more likely to rely on screens at age 3, suggesting that early vulnerabilities amplify later screen use, highlighting the sensitivity of language networks to environmental input and the potential for screen exposure to alter developmental trajectories.

---

Language is distinct from other developmental abilities such as motor or cognitive skills in that it is typically lateralized to the left hemisphere, although it begins bilaterally represented in early life.^1^ Language specialization may be more dependent on reciprocal interaction and environmental inputs, unlike motor skills, which emerge more robustly. In turn, toddlerhood represents a critical period of language development, supported by both neurobiological maturation and environmental experiences.^1,2^ The brain’s language regions undergo rapid myelination during this stage,^3,4^ which aligns with reported structural and functional changes such as increased cortical thickness and interhemispheric connectivity between Broca’s area and its right homologue.^5,6^

As language development requires reciprocal interaction, replacing input with passive or incoherent noise (e.g., TV, excessive screen exposure) risks disrupting this sensitive developmental trajectory. Converging evidence now indicates that early and excessive screen time exposure can negatively impact children’s neurodevelopment^7^, particularly in language domains.^8,9^ However, these associations may be bidirectional – delayed language may increase dependency on screens, reducing demands on reciprocal communication but offering immediate and reinforcing stimulation.^10^

In the current study, we examined associations between screen time, volumes of speech-language centres, and language outcomes at age 2 years, as well as their longitudinal associations from ages 2 to 3 years. We focused on two key questions: [R1] Do regional brain volumes mediate the association between screen time and language outcomes at age 2? [R2] Do regional brain volumes and language outcomes at age 2 predict screen time use at age 3? We hypothesized that more time spent on screens at age 2 would be associated with smaller volumes in speech– language regions and poorer concurrent language outcomes, and that these effects would prospectively predict greater reliance on screens at age 3. As part of an exploratory analyses, we further examined whether screen time at age 2 was associated with pars triangularis volumes at age 3.

Toddlers (N=70) were from the *Pregnancy during the COVID-19 Pandemic* cohort (Supplementary Table 1). Mothers reported screen time hours on typical weekdays and weekends at 2 and 3 years, and 2-year-old language outcomes using the MacArthur–Bates Communicative Development Inventories (CDI). Toddlers completed magnetic resonance imaging at age 2 (N=70), with a subset at age 3 years (N=32). T1-weighted images were processed using Infant FreeSurfer to extract 68 cortical and 20 subcortical volumes, normalized by total cerebral volume (Supplementary Information).

To identify brain regions associated with both screen time and language scores, we examined the correlation coefficients with each measure with regional brain volumes at age 2. Smaller right pars triangularis volumes were associated with increased screen time (r=-.26, p=.03), while larger volumes in this region were associated with higher language scores (CDI, r=.27, p=.03) (Supplementary Table 2).

Structural equation modeling first tested whether right pars triangularis volumes mediated the association between screen time and 2-year CDI scores. A second longitudinal mediation model examined whether the effect of 2-year right pars triangularis volumes on 3-year screen time was mediated by 2-year language outcomes.

Exploratory regression analyses examined whether screen time at age 2 predicted right pars triangularis volumes at age 3.

At age 2, children’s average screen time was 0.98±1.25hr on weekdays and 1.28±1.25hrs on weekends. By age 3, weekday screen time remained similar (1.04±1.03hr), while weekend screen time increased to 1.78±1.40hr.

In the cross-sectional mediating model (χ^2^(1)=0.95, p=.33, CFI=1.00, TLI=1.00, RMSEA=.00), higher screen time at age 2 significantly predicted smaller right pars triangularis volumes (β=−.26, p=.037) and poorer 2-year language outcomes (β=−.40, p=.001). However, the volumes did not predict language outcomes (β=.15, p=.17; Figure 1A), indicating that the effects of screen time on brain development and language appeared to be occurring in parallel rather than mediated.

**Figure 1A.**
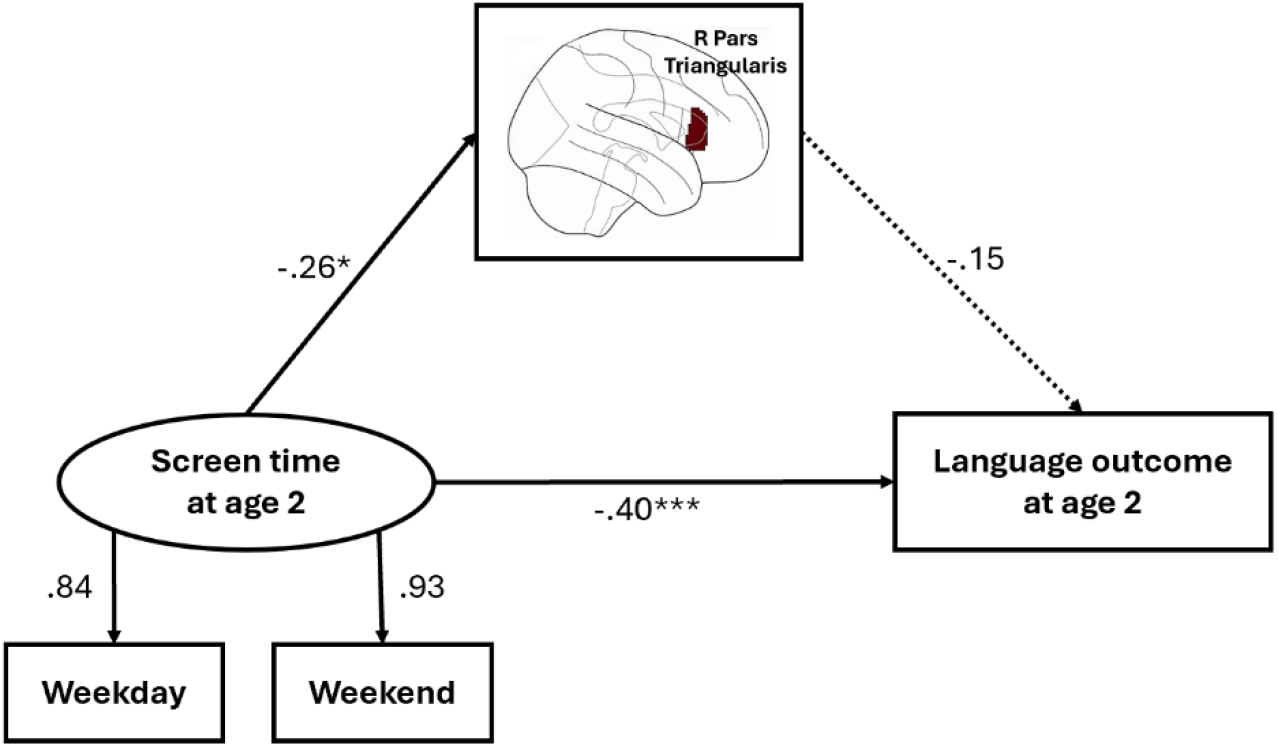
Associations between screen time, regional brain volume, and language outcome at age 2

In the second longitudinal mediation model, with 2 and 3 year data, (χ^2^(1)=0.51, p=.47, CFI=1.00, TLI=1.00, RMSEA=.00), smaller right pars triangularis volumes at age 2 predicted longer ST use at age 3 (β=−.42, p=.002), fully through poorer language outcomes at age 2 (β=.26, p=.026; Figure 1B).

Exploratory analyses showed that weekday screen time at age 2 was positively associated with left pars triangularis volumes at age 3 (β=.71, p=.034; Figure 1C), but not right pars triangularis volumes (weekday screen time: β=.09, p=.794; weekend screen time: β=.01, p=.987).

Early in development, screen time showed parallel effects on brain structure and language ability, with higher exposure linked to both smaller pars triangularis volumes and poorer language outcomes at age 2. Over time, however, children with smaller volumes and weaker language skills at age 2 became more likely to rely on screens at age 3, suggesting that early vulnerabilities amplify later screen use. Given that toddlerhood represents an important period for language development and hemispheric specialization, these findings highlight the sensitivity of language networks to environmental input and the potential for screen exposure to alter developmental trajectories.

Exploratory analyses further indicated that screen time at age 2 predicted larger left pars triangularis volumes at age 3, but not right-sided volumes. This shift from right to left hemisphere involvement may reflect accelerated or compensatory lateralization in response to screen-based input, raising the possibility that early screen exposure alters the trajectory of language development.

Findings underscore the potential for targeted interventions during toddlerhood to support vulnerable groups. The possible long-term effects of excessive screen exposure on language lateralization warrant further investigation.

**Figure 1B.**
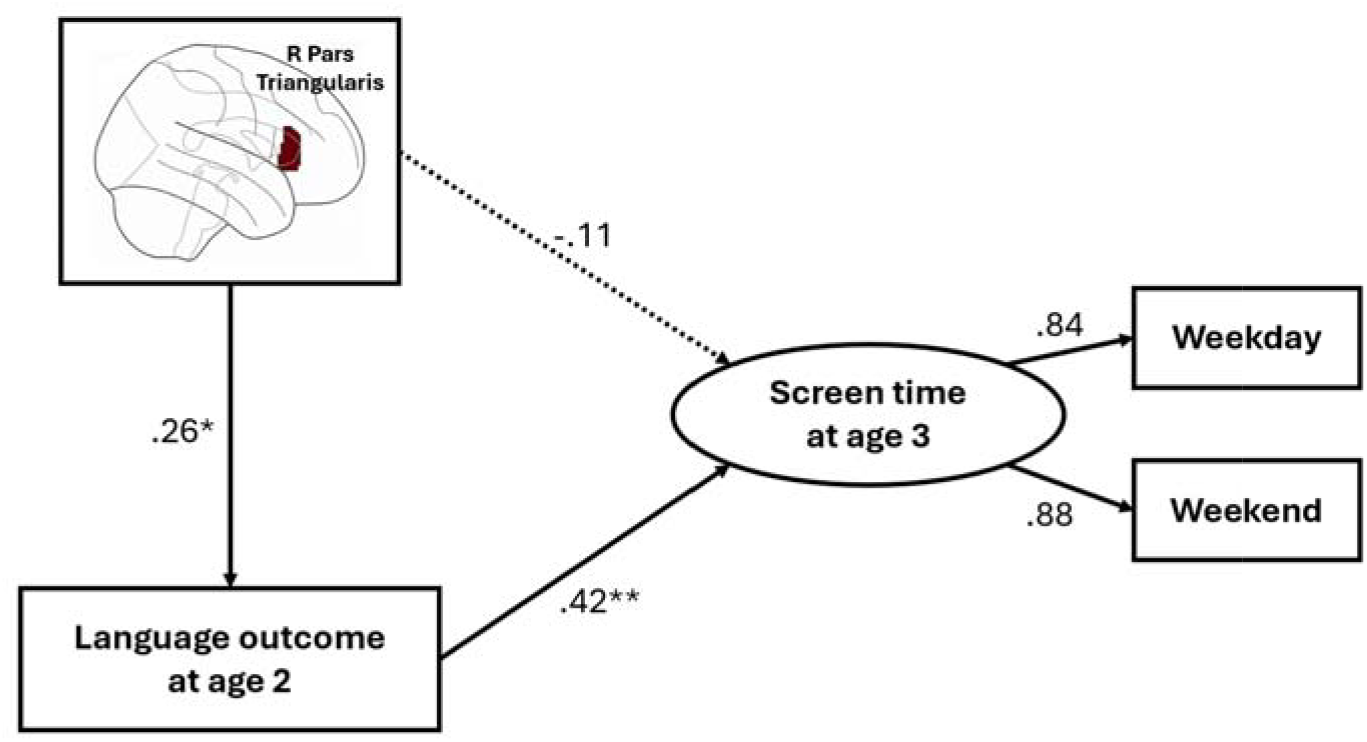
Longitudinal effect of regional brain volume and language outcome at age 2 on screen time at age 3

**Figure 1C.**
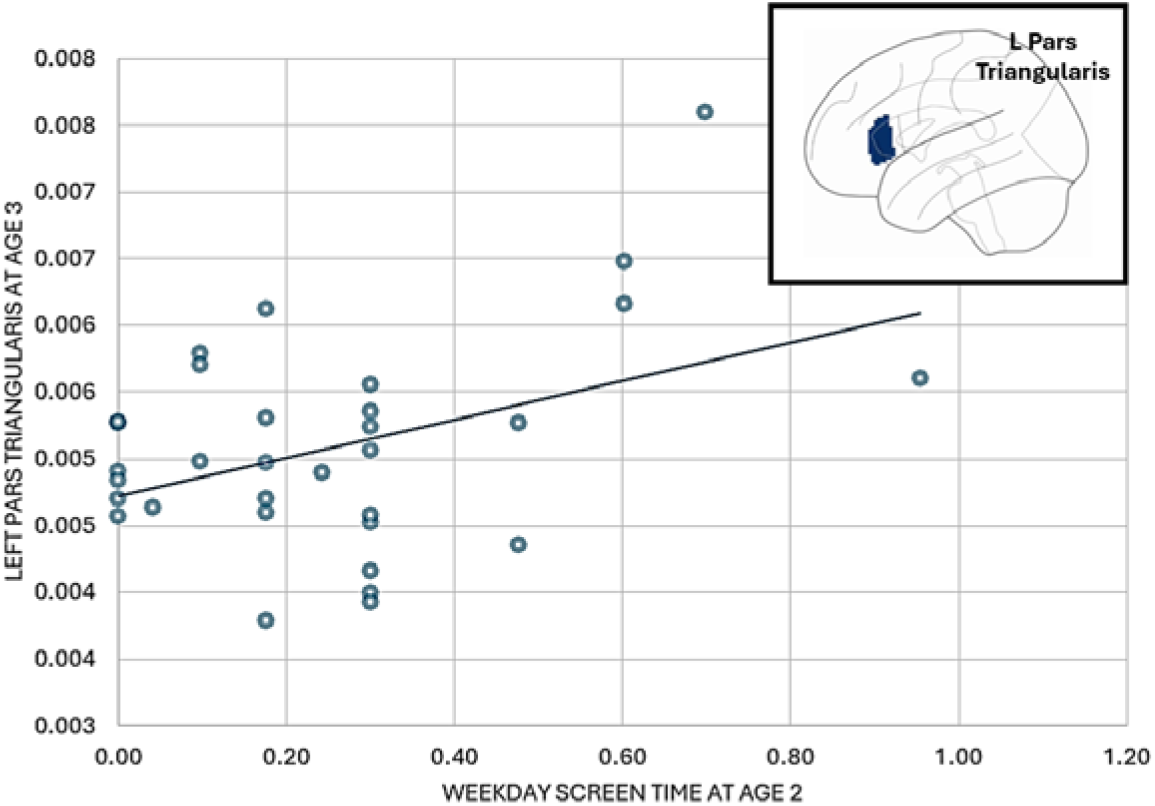
The associations between screen time at age 2 and left pars triangularis volume at age 3

## Acknowledgements

This project would have not been possible without the participation from all families in the PdP study and the authors extend their sincerest thanks. This work was supported by the Alberta Children’s Hospital Research Institute, the Owerko Centre for Neurodevelopment and Mental Health, and the Canadian Institutes of Health Research (UIP 178826). CL, EGD, LT-M receive funding from the Canada Research Chairs Program.

