## Supplemental information for "Longitudinal Associations Between Screen Time, Brain Development, and Language Outcomes in Early Childhood"

**Supplementary Information. T1-weighted image acquisition and preprocessing**

Image Acquisition: infants were scanned using a GE 3T MR750w MRI with a 32-channel head coil at the Alberta Children’s Hospital to acquire brain imaging data. All infants were scanned while asleep atop an inflatable MedVac infant scanning bed. T1-weighted images were obtained (repetition time = 5200 ms, echo time = 2200 ms, inversion time = 540 ms, field of view = 1900 mm, matrix = 512 X 512, bandwidth = 41.67, voxel 1 X 1 X 1 mm^3, flip angle = 12°, 136 slices, total time = 3:32).

Preprocssing: Cortical reconstruction and volumetric segmentation of all neonatal, infant, and toddler data was performed using Infant FreeSurfer^1–3^, an automated segmentation and surface extraction pipeline for infant T1-weighted neuroimaging data, which is documented and freely available for download online (https://surfer.nmr.mgh.harvard.edu/fswiki/infantFS). The technical details of these procedures are described in prior publications^4–16^. Briefly, this processing includes motion correction and averaging^15^ of multiple volumetric T1 weighted images (when more than one is available), removal of non-brain tissue using a hybrid watershed/surface deformation procedure^14^, automated Talairach transformation, segmentation of the subcortical white matter and deep gray matter volumetric structures (including hippocampus, amygdala, caudate, putamen, ventricles)^7,8^, intensity normalization^17^, tessellation of the gray matter white matter boundary, automated topology correction^6,18^, and surface deformation following intensity gradients to optimally place the gray/white and gray/cerebrospinal fluid borders at the location where the greatest shift in intensity defines the transition to the other tissue class^5,19^. Once the cortical models are complete, a number of deformable procedures can be performed for further data processing and analysis including surface inflation^9^, registration to a spherical atlas which is based on individual cortical folding patterns to match cortical geometry across subjects^10^, parcellation of the cerebral cortex into units with respect to gyral and sulcal structure^11,20^, and creation of a variety of surface based data including maps of curvature and sulcal depth. This method uses both intensity and continuity information from the entire three-dimensional MR volume in segmentation and deformation procedures to produce representations of cortical thickness, calculated as the closest distance from the gray/white boundary to the gray/CSF boundary at each vertex on the tessellated surface^5^. The maps are created using spatial intensity gradients across tissue classes and are therefore not simply reliant on absolute signal intensity. The maps produced are not restricted to the voxel resolution of the original data and thus can detect submillimeter differences between groups. Procedures for the measurement of cortical thickness have been validated against histological^21^ and manual measurements^22,23^. Freesurfer morphometric procedures have been demonstrated to show good test-retest reliability across scanner manufacturers and across field strengths^12,16^. Age in months is specified when running Infant Freesurfer, and the segmentation atlas is built from the four participants closest in age. All segmentations were examined by a trained research assistant to ensure accuracy.

**Supplementary Table 1. Demographic information**

| Variable | | Mean ± SD / N (%) |
| --- | --- | --- |
| Gestational age at birth (weeks, Mean ± SD) | | 39.25 ± 1.18 |
| Chid birth weight (gram, Mean ± SD) | | 3358.61 ± 412. 85 |
| Mother’s age (year, Mean ± SD) | | 33.22 ± 3.71 |
| Maternal marital status (N, %) | Single | 1 (1.4%) |
|  | Married | 59 (84.3%) |
|  | Common-law | 9 (12.9%) |
|  | Separated | 1 (1.4%) |
| Household income (N, %) | Less than $69,999 | 5 (7.2%) |
|  | $70,000 to $99,999 | 13 (18.6%) |
|  | $100,000 to $149,999 | 15 (21.5%) |
|  | $150,000 to $199,999 | 17 (24.3%) |
|  | Over $200,000 | 20 (28.6%) |
| Maternal education (N, %) | Completed high school | 3 (4.3%) |
|  | Completed college | 11 (15.7%) |
|  | Bachelor's Degree | 33 (47.1%) |
|  | Master's Degree | 14 (20.0%) |
|  | Doctorate | 9 (12.9%) |

**Supplementary Table 2. Correlation matrix between screen time and regional brain volumes**

|  | Weekday screen time | Weekend screen time | Langugage outcome |
| --- | --- | --- | --- |
| bankssts_L | -0.111 | -0.020 | 0.054 |
| caudalanteriorcingulate_L | -0.069 | -0.114 | -0.025 |
| caudalmiddlefrontal_L | .291* | .303* | -0.122 |
| cuneus_L | 0.102 | 0.018 | 0.158 |
| entorhinal_L | 0.071 | 0.067 | 0.084 |
| fusiform_L | -0.010 | 0.086 | 0.010 |
| inferiorparietal_L | -0.009 | 0.070 | -0.144 |
| inferiortemporal_L | 0.075 | 0.194 | -0.067 |
| isthmuscingulate_L | 0.055 | 0.050 | 0.025 |
| lateraloccipital_L | 0.057 | 0.063 | 0.177 |
| lateralorbitofrontal_L | 0.041 | 0.105 | -0.046 |
| lingual_L | -0.125 | -0.081 | 0.194 |
| medialorbitofrontal_L | -0.069 | -0.034 | -0.125 |
| middletemporal_L | -0.215 | -0.127 | 0.106 |
| parahippocampal_L | -0.119 | -0.073 | .352** |
| paracentral_L | 0.092 | 0.022 | -.279* |
| parsopercularis_L | 0.099 | 0.115 | 0.075 |
| parsorbitalis_L | -0.141 | -0.113 | -0.093 |
| parstriangularis_L | -0.074 | -0.100 | 0.112 |
| pericalcarine_L | -0.026 | -0.064 | .261* |
| postcentral_L | -0.131 | 0.014 | 0.111 |
| posteriorcingulate_L | -0.180 | -0.158 | -0.024 |
| precentral_L | -0.074 | 0.006 | -0.012 |
| precuneus_L | -0.083 | -0.172 | -0.035 |
| rostralanteriorcingulate_L | -0.034 | 0.025 | 0.057 |
| rostralmiddlefrontal_L | -0.216 | -.297* | 0.038 |
| superiorfrontal_L | 0.125 | 0.069 | -0.128 |
| superiorparietal_L | 0.041 | -0.053 | 0.150 |
| superiortemporal_L | -0.121 | 0.014 | -0.110 |
| supramarginal_L | -0.135 | -0.068 | -0.054 |
| frontalpole_L | 0.035 | 0.135 | -0.044 |
| temporalpole_L | 0.016 | 0.061 | -0.216 |
| transversetemporal_L | 0.123 | 0.029 | -0.085 |
| insula_L | -0.023 | 0.006 | 0.155 |
| bankssts_R | -0.007 | -0.037 | 0.021 |
| caudalanteriorcingulate_R | -0.167 | -0.091 | -0.021 |
| caudalmiddlefrontal_R | .249* | 0.191 | -0.076 |
| cuneus_R | -0.064 | -0.039 | 0.147 |
| entorhinal_R | 0.011 | -0.057 | 0.150 |
| fusiform_R | -0.042 | -0.035 | 0.072 |
| inferiorparietal_R | -0.046 | -0.044 | 0.071 |
| inferiortemporal_R | 0.003 | 0.043 | -0.021 |
| isthmuscingulate_R | -0.069 | -0.054 | .260* |
| lateraloccipital_R | 0.100 | 0.138 | 0.138 |
| lateralorbitofrontal_R | 0.029 | 0.043 | -0.063 |
| lingual_R | -0.013 | 0.045 | 0.080 |
| medialorbitofrontal_R | -0.098 | -0.092 | 0.012 |
| middletemporal_R | 0.040 | 0.168 | -0.093 |
| parahippocampal_R | -0.054 | 0.033 | 0.227 |
| paracentral_R | -0.170 | -0.212 | 0.066 |
| parsopercularis_R | 0.085 | 0.084 | 0.033 |
| parsorbitalis_R | -0.026 | -0.168 | -0.013 |
| parstriangularis_R | -0.176 | -.262* | .266* |
| pericalcarine_R | 0.018 | 0.050 | 0.128 |
| postcentral_R | 0.066 | 0.171 | -0.206 |
| posteriorcingulate_R | -0.154 | -0.162 | 0.126 |
| precentral_R | 0.175 | 0.234 | -0.157 |
| precuneus_R | -0.094 | -0.060 | 0.026 |
| rostralanteriorcingulate_R | -0.224 | -0.116 | 0.028 |
| rostralmiddlefrontal_R | -0.076 | -0.193 | 0.084 |
| superiorfrontal_R | -0.103 | 0.004 | 0.074 |
| superiorparietal_R | -0.055 | 0.076 | -0.020 |
| superiortemporal_R | 0.119 | .278* | -0.169 |
| supramarginal_R | 0.043 | 0.026 | 0.010 |
| frontalpole_R | -0.119 | -0.213 | -0.079 |
| temporalpole_R | 0.112 | 0.146 | -0.035 |
| transversetemporal_R | 0.170 | 0.104 | -0.223 |
| insula_R | -0.083 | -0.189 | 0.195 |
| Thalamus_L | -0.008 | -0.066 | 0.005 |
| Caudate_L | 0.078 | 0.213 | -.253* |
| Putamen_L | -0.057 | -0.116 | 0.035 |
| Pallidum_L | 0.077 | 0.016 | -0.044 |
| Hippocampus_L | 0.084 | 0.169 | -.260* |
| Amygdala_L | -0.187 | -0.067 | -0.160 |
| Accumbens_area_L | -0.013 | 0.109 | -0.122 |
| VentralDC_L | 0.140 | 0.177 | -0.212 |
| Thalamus_R | 0.052 | 0.001 | -0.135 |
| Caudate_R | -0.056 | -0.036 | 0.006 |
| Putamen_R | -0.031 | 0.005 | -0.035 |
| Pallidum_R | 0.193 | 0.205 | -.253* |
| Hippocampus_R | -0.059 | 0.123 | -0.050 |
| Amygdala_R | -0.048 | 0.019 | -0.049 |
| Accumbens_area_R | 0.069 | 0.116 | -0.054 |
| VentralDC_R | 0.061 | 0.132 | -0.213 |
| Vermis | 0.057 | 0.047 | 0.092 |
| Midbrain | 0.051 | 0.033 | 0.004 |
| Pons | 0.037 | 0.070 | -.242* |
| Medulla | 0.128 | 0.157 | 0.045 |
